# Rescuing Photoreceptors in RPE Dysfunction-Driven Retinal Degeneration: The Role of Small Extracellular Vesicles Secreted from Retinal Pigment Epithelium

**DOI:** 10.1101/2024.04.09.588773

**Authors:** Dimitrios Pollalis, Constantin Georgescu, Jonathan D. Wren, Grigor Tombulyan, Justin M. Leung, Pein-An Lo, Clarisa Marie Bloemhof, Ryang Hwa Lee, EunHye Bae, Jeffrey K. Bailey, Britney O. Pennington, Amir I. Khan, Kaitlin R. Kelly, Ashley K. Yeh, Kartik S. Sundaram, Mark Humayun, Stain Louie, Dennis O. Clegg, Sun Young Lee

**Affiliations:** Roski Eye Institute, Keck School of Medicine, University of Southern California, Los Angeles, CA, USA; Genes & Human Diseases Research Program, Oklahoma Medical Research Foundation, Oklahoma City, USA; Department of Cell Biology and Genetics, Institute for Regenerative Medicine, School of Medicine, Texas A&M University, TX, USA; Center for Stem Cell Biology and Engineering, Neuroscience Research Institute, University of California, Santa Barbara, CA, USA; Mann School of Pharmacy and Pharmaceutical Sciences, University of Southern California, Los Angeles, CA, USA; Department of Physiology and Neuroscience, Keck School of Medicine, University of Southern California, USA

## Abstract

Dysfunction of the retinal pigment epithelium (RPE) is a common shared pathology in major degenerative retinal diseases despite variations in the primary etiologies of each disease. Due to their demanding and indispensable functional roles throughout the lifetime, RPE cells are vulnerable to genetic predisposition, external stress, and aging processes. Building upon recent advancements in stem cell technology for differentiating healthy RPE cells and recognizing the significant roles of small extracellular vesicles (sEV) in cellular paracrine and autocrine actions, we investigated the hypothesis that the RPE-secreted sEV alone can restore essential RPE functions and rescue photoreceptors in RPE dysfunction-driven retinal degeneration. Our findings support the rationale for developing intravitreal treatment of sEV. We demonstrate that intravitreally delivered sEV effectively penetrate the full thickness of the retina. Xenogenic intraocular administration of human-derived EVs did not induce acute immune reactions in rodents. sEV derived from human embryonic stem cell (hESC)-derived fully differentiated RPE cells, but not sEV-depleted conditioned cell culture media (CCM minus sEV), rescued photoreceptors and their function in a Royal College of Surgeons (RCS) rat model. This model is characterized by photoreceptor death and retinal degeneration resulting from a mutation in the *MerTK* gene in RPE cells. From the bulk RNA sequencing study, we identified 447 differently expressed genes in the retina after hESC-RPE-sEV treatment compared with the untreated control. Furthermore, 394 out of 447 genes (88%) showed a reversal in expression toward the healthy state in Long-Evans (LE) rats after treatment compared to the diseased state. Particularly, detrimental alterations in gene expression in RCS rats, including essential RPE functions such as phototransduction, vitamin A metabolism, and lipid metabolism were partially reversed. Defective photoreceptor outer segment engulfment due to intrinsic *MerTK* mutation was partially ameliorated. These findings suggest that RPE-secreted sEV may play a functional role similar to that of RPE cells. Our study justifies further exploration to fully unlock future therapeutic interventions with sEV in a broad array of degenerative retinal diseases.

## INTRODUCTION

Retina pigment epithelium (RPE) cells, uniquely present in the retina located between the photoreceptors and the choroid vasculature, serves highly specialized functions. These include phagocytosis of photoreceptor outer segments (POS), conservation of phototransduction and visual cycle, protection against oxidative stresses, secretion of cytokines and growth factors, and transportation of nutrients or ions.**1-4** Due to their demanding and indispensable roles they play throughout the lifetime, RPE cells are vulnerable to genetic predisposition, external stress, and aging processes. **4-6** Therefore, dysfunctional RPE cells commonly represent a shared pathology in major retinal degenerative conditions such as age-related macular degeneration (AMD) and inherited retinal diseases (IRDs). While the precise pathobiology may vary among different retinal diseases, it is widely acknowledged that progressive RPE dysfunction, occurring slowly and heterogeneously, precedes RPE cell death. This, in turn, leads to irreversible damage to photoreceptors due to the impaired symbiotic relationship between RPE and photoreceptor, which is a recurring feature in many degenerative retinal conditions. **4-7**

Of the various strategies under investigation to address RPE cell dysfunction as an approach for treating retinal degenerative conditions, the RPE secretome holds great potential, benefiting from technological advancements in differentiating healthy RPE cells from pluripotent stem cells (PSC) or induced PSC (iPSC). These fully differentiated and polarized RPE cells have proven intraocular safety profiles from ongoing human clinical trials. **8-11** The strength of the RPE secretome lies in employing a multimolecular approach, better supporting the multifunctional nature of the RPE, whereas an approach targeting a single molecule is often suboptimal in therapeutic efficacy.

While the composition of the secretome is dynamic, depending on the cell type and microenvironmental stimuli, generally, the stem cell secretome is known to have therapeutic benefits for tissue repair including proangiogenic, antiapoptotic, antifibrotic, anti-inflammatory, and immunomodulatory effects. Thus, previous research on the RPE secretome, including our previous study, has primarily focused on trophic factors and their role in neuroprotection. **3, 12-13** However, accumulating evidence suggests that small extracellular vesicles (sEV) play a crucial role in mediating cell-to-cell communication. **14-17** In our recent study, we investigated the dynamics of microRNA (miRNA) expression contained within sEV secreted during the maturation of RPE cells after their full differentiation derived from human embryonic stem cells (hESC). We examined three stages of maturation: early stage (11-22 days in culture), mid-stage (28-39 days in culture), and late stage (59-70 days in culture) hESC-derived RPE cells. Pathway analyses of miRNA from mid-stage hESC-RPE cells, compared to other time points, revealed significant involvement in key RPE functions such as tissue and cell polarity, response to growth factors, inflammation, oxidative stress, and senescence. These findings suggest that the cargo sorting, for example miRNA, of RPE cell-derived sEV is tightly regulated and reflects the functional state of the sEV secreting cells. **18** This provides a rationale to prioritize testing the therapeutic efficacy of mid-stage (28-39 days in culture) hESC-RPE secreted sEV.

In the present study, we examined our hypothesis that the RPE-secreated sEV alone can restore essential RPE functions and rescue photoreceptors in RPE dysfunction-driven retinal degeneration. We investigated whether sEV released from fully differentiated hESC-RPE could effectively rescue photoreceptors in the Royal College of Surgeons (RCS) rat model, animal models of retinal degeneration characterized by PR death due to a mutation in the *MerTK* gene in RPE. Additionally, we explored the molecular pathways associated with the therapeutic effects of hESC-RPE treatment, with a special focus on essential RPE functions.

## RESULTS

### Characterization of fully differentiated RPE cells from human embryogenetic stem cell

The cultured RPE cells fully differentiated from hESC-RPE displayed a characteristic cobblestone appearance with hexagonal shapes and intermediate pigmentation levels (**Figure 1-A**). The Immunocytochemistry examination confirmed the presence of RPE-specific markers with a distinct staining pattern, utilizing antibodies against BEST1, RPE65, and ZO-1 proteins (**Figure 1-B**).

**Figure 1.**
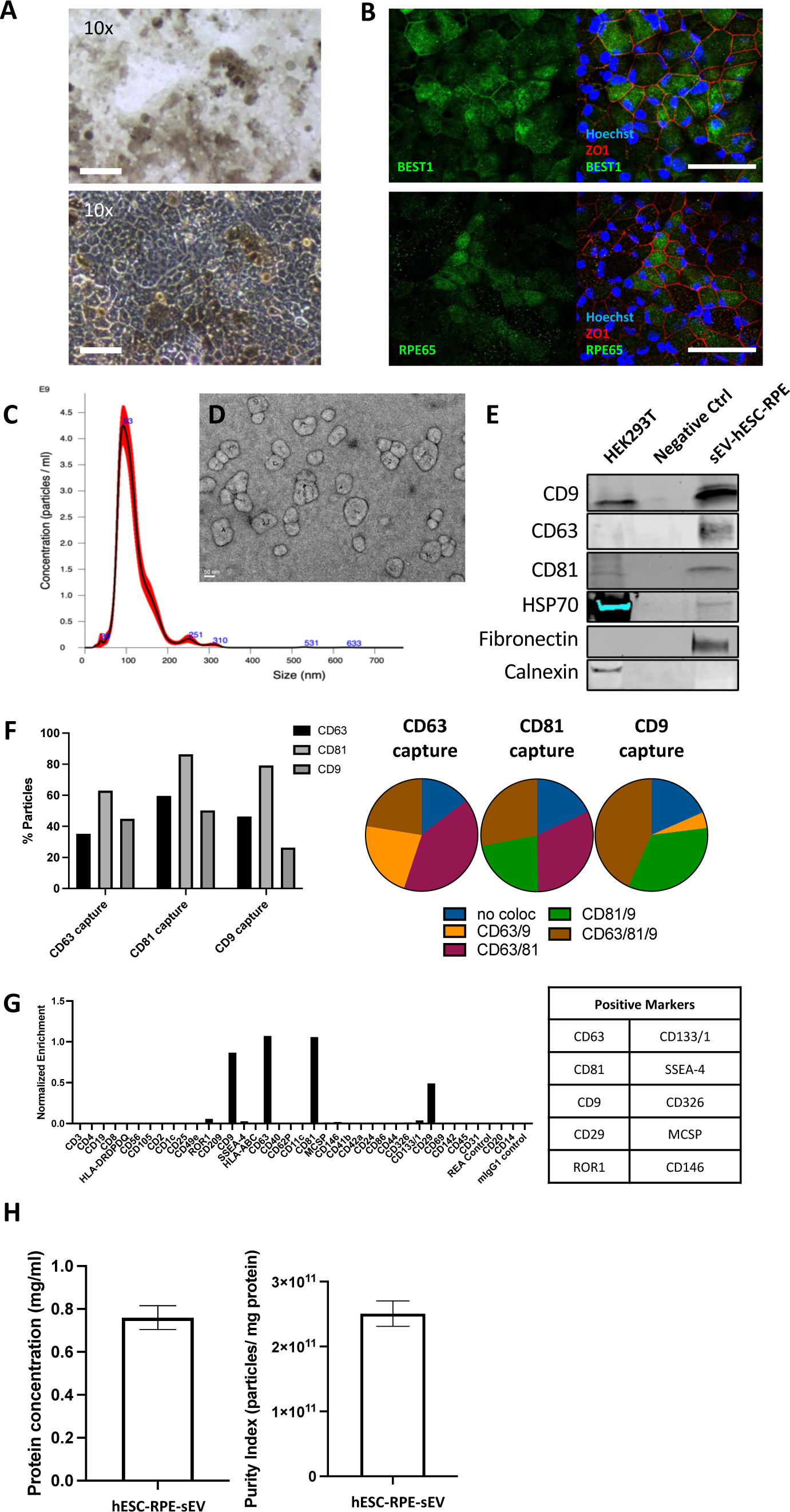
Characterization of human embryogenic stem cell derived fully differentiated retina pigment epithelium and their secreted small extracellular vesicles. **A**. Representative brightfield and phase-contrast microscopy images demonstrating pigmented hexagonal hESC-RPE cells exhibiting cobblestone morphology (scale bar, 100 μm). **B**. Immunofluorescence images depicting hESC-RPE cells stained with anti-BEST1, RPE65, and ZO1 antibodies, highlighting RPE cellular markers (scale bar, 50 μm). **C**. Size-concentration graph illustrating sEV derived from hESC-RPE cells, analyzed via NanoSight, revealing a count of 8.56 x 10^10^ (+/- 1.6 x 10^10^) per mL and an average size of 93 nm.**D**. Representative TEM image revealing the characteristic morphology of sEV. **E**. Western blot analysis confirming the presence of EV markers including CD9, CD63, CD81, and HSP70. **F.** Distribution and colocalization analysis of tetraspanins (CD81, CD9, and CD63) within hESC-RPE-sEV. **G**. MACSPlex assay results indicating positivity for extended surface markers (CD63, CD81, CD9, CD29, ROR1, CD133/1, SSEA-4, CD326, MCSP, CD146) in hESC-RPE-sEV. **H**. Graphs presenting protein concentration and purity index (particle-to-protein ratio) of the recovered hESC-RPE-sEV.

### Biophysical and molecular characterization of hESC-RPE secreted sEV

The quantified sEV particle numbers were 8.56 x 10^10^ (+/- 1.6 x 10^10^) per mL, with an average size of 93 nm (**Figure 1-C**). Characteristic sEV morphology, showing a prominent cup shape and intact bilipid membrane structures, was confirmed by Transmission Electron Microscopy (TEM) (**Figure 1-D**). Western blot analysis validated the presence of EV markers, including CD9, CD63, CD81 and HSP70 in **Figure 1-E**. The distribution of tetraspanins, determined by ExoView analysis, is illustrated in **Figure 1-F**. Additionally, surface protein identification using 37 surface marker antibodies through the MACSPlex assay identified CD63, CD133/1, CD81, SSEA-4, CD9, CD326, CD29, MCSP, ROR1, and CD146 (**Figure 1-G**).

The protein concentration in hESC-RPE-sEV was determined to be 0.76 mg/mL with the purity index, indicated by the particle-to-protein ratio of the recovered sEV being 2.51 x 10^10^ (**Figure 1-H**).

### Intravitreally delivered hESC-RPE-sEV reach full-thickness retina

Despite the limitation in detecting individual particles within the retina, confocal microscopy analysis of retinal sections obtained 24 hours following the intravitreal injection of fluorescently tagged hESC-RPE-sEV in Long-Evans (LE) rats revealed their presence in both the inner and outer retina, indicating their penetrating through the full thickness in a scattered manner (**Figure 2-A**).

**Figure 2.**
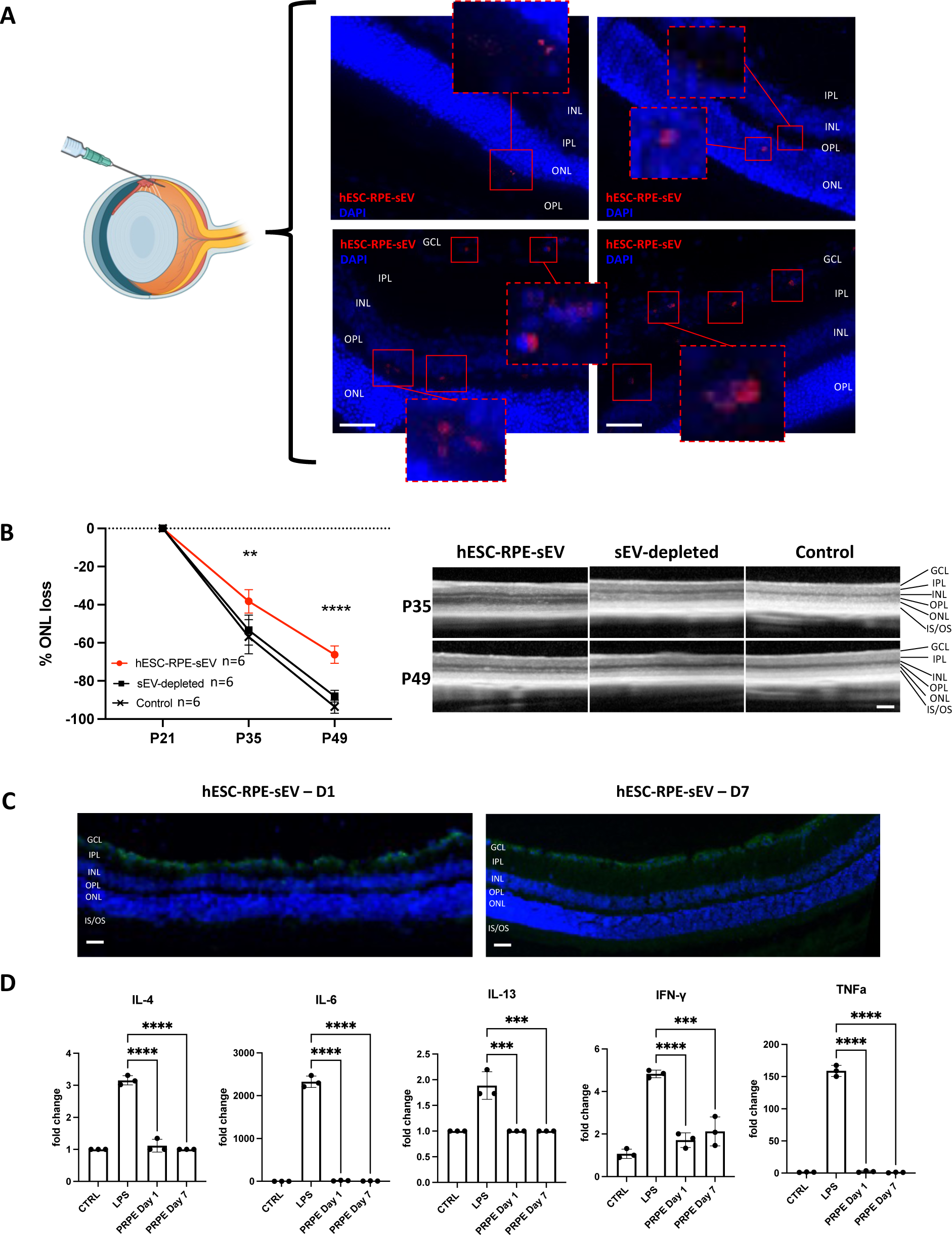
Assessing retinal uptake xenogenic intraocular immunoreaction of intravitreally delivered hESC-RPE-sEV and bioactive fraction of CCM from hES-sEV. A. Intravitreally delivered hESC-RPE-sEV reach the full thickness of the retina. Retinal sections, obtained 24 hours post-injection in Long-Evans (LE) rats, demonstrate the distribution of fluorescently (MemGLowTM 640) labeled hESC-RPE-sEV (*red*) within the retina, displaying penetration into both inner and outer retinal layers in a scattered pattern (scale bar, 50 μm). **B. The sEV fraction, not the sEV-depleted fraction, of hESV-RPE cell-conditioned medium (CCM), preserved photoreceptors in immunocompromised RCS rats**. Immunocompromised RCS rats (iRCS) received two intravitreal injections of either sEV fraction (∼ 5 x 10^8^/10 μl) or sEV depleted fraction (undetectable particles/10 μl) of hESC-RPE-CCM at P21 and P35. *In vivo* Optical coherence tomography (OCT) analysis showed a significant preservation of outer nuclear layer (ONL) thickness in eyes treated with hESC-RPE-sEV (18.61%; *p-value 0.002* and and 27.31%; *p-value < 0.0001* respectively; *n*=6 in each group) compared to sEV-depleted sample-treated and untreated control eyes at P35 and P49. (scale bar, 200 μm)**. C and D**. **Intravitreally delivered xenogenic hESC-RPE-sEV did not induce reactive retinal gliosis or acute immune reaction.** A single intravitreal injection of hESC-RPE-sEV was given to Long-Evans (LE) Rats (∼ 5 x 10^8^/10 μl) and C57BL/6 mic (∼ 5 x 10^7^/1 μl). **C**. Representative immunofluorescence images of GFAP staining in LE rat retinas at 1 and 7 days post treatment, illustrating the absence of reactive gliosis in addition to base line GFAP expressions. **D**. PCR analysis showed no increase in acute inflammation markers (IL-4, IL-6, IL-13, IFN-γ, and TNF-α) after treatment in C57BL/6J mice compared to untreated controls (*n*=3 in each group).

### hESC-RPE-sEV treatment, not sEV depleted conditioned cell culture media (CCM), rescues photoreceptor in immunocompromised RCS rats

To assess the varying efficacy of different fractions of CCM from hESC-RPE between hESC-RPE-sEV and sEV-depeleted fraction (CCM minus sEV), immunocompromised RCS (iRCS) rats received IVT injections with the sEV-enriched or sEV-depleted fractions at P21 and P35. Eyes injected with hESC-RPE-sEV exhibited an 18.61% (*p-value 0.002*) and 27.31% (*p-value < 0.0001*) preservation of the outer nuclear layer (ONL) compared to the control (no injection) groups at P35 and P49 respectively, as assessed by OCT imaging. Importantly, the sEV-depleted fraction treatment group did not show photoreceptor preservation compared to the control (**Figure 2-B**).

### Xenogenic intraocular hESC-RPE-sEVs treatment do not induce acute immune reaction

We confirmed that IVT injection of hESC-RPE-sEV did not induce immediate reactive gliosis in non-degenerative retinas of LE rats at 1 and 7 days post-injection. This suggests that treatment with xenogenic intraocular hESC-RPE-sEV did not prompt immediate reactive retinal gliosis in LE rats (**Figure 2-C**). Wild-type (C57BL/6) mice received IVT injections of either hESC-RPE-sEV or Lipopolysaccharides (LPS) (1 ng/μL) and were evaluated at 1- and 7-days post-injection. PCR analysis for IL-4, IL-6, IL-13, IFN-γ, and TNF-α revealed no increase in inflammation markers in the eyes injected with hESC-RPE-sEV (*n*=3). These results were comparable to the untreated control group and significantly different from the LPS-injected group (**Figure 2-D**).

### hES-RPE-sEV treatment rescue photoreceptors and theirs function in RCS rats

Following two IVT injections at P21 and P35, the progressive loss of outer nuclear layer (ONL) was significantly rescued by 11% (*p-value 0.001*) at P49 in the hESC-RPE-sEV treated group (*n*=18) (**Figure 3-A**). The functional preservation of the anatomically preserved ONL in OCT was assessed using ERG. ERG examination revealed significantly preserved scotopic *b*-wave amplitudes in the hESC-RPE-sEV treated group with a similar trend in scotopic *a*-wave amplitude (*n*=17) (**Figure 3-B**). Photopic *a*- and *b*-wave amplitudes showed no significant difference between the treated and untreated groups.

**Figure 3.**
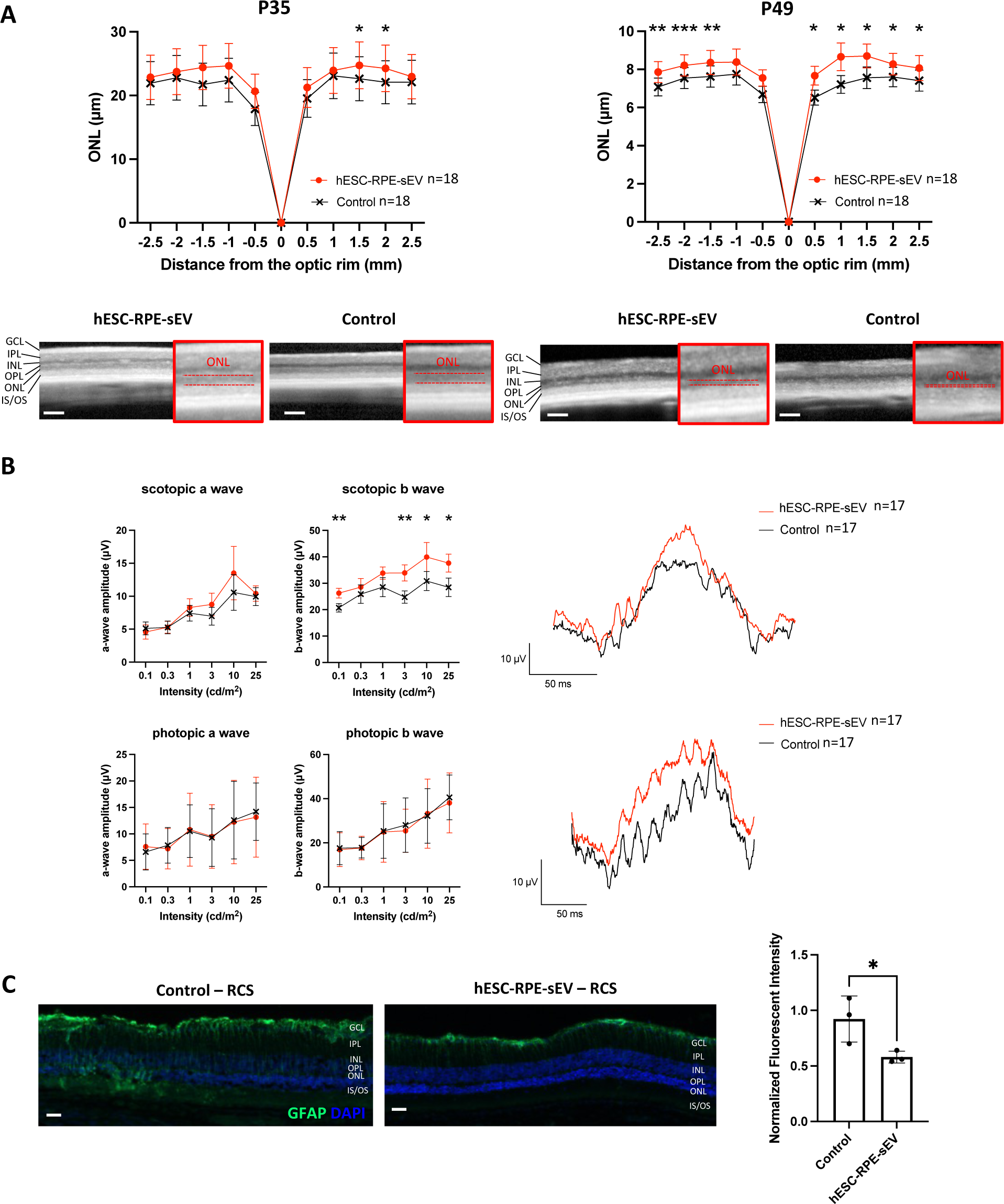
Intravitreal injections of hESC-RPE-sEV treatment in RCS rats rescue photoreceptor and their function. Two consecutive intravitreal injection of hESC-RPE-sEV (∼ 5 x 10^8^/10 μl) were administered to one eye of RCS Rats at P21 and P35. **A.** OCT analysis showed a significant difference in outer nuclear layer (ONL) thickness preservation in eyes treated with hESC-RPE-sEV compared to untreated control eyes at P35 and P49. Representative OCT images at P35 and P49 are provided. (*n*=18, scale bar, 200 μm). **B**. Electroretinography (ERG) analysis (*n*=17). **C**. Immunofluorescence images of GFAP staining in RCS rat retinas at P49 (*n*=3). * p-value < 0.05, ** p-value < 0.01, *** p-value < 0.001.

hESC-RPE-sEV treatment also reduced the abnormally elevated retinal glial activation observed during the natural course of retinal degeneration in RCS rats by 36.46% (*p-value 0.05*). This reduction was semi-quantified by GFAP expression levels in comparison to the untreated control group (**Figure 3-C**).

### Transcriptomic analysis in RCS rats followed by hESC-RPE-sEV treatment

To investigate the underlying mechanism responsible for rescuing photoreceptors in RCS rats treated with hESC-RPE-sEV, we conducted mRNA-seq analysis to identify differentially expressed genes (DEGs) in RCS retinas with and without hESC-RPE-sEV treatment. The analysis identified a total of 447 DEGs (with a fold change FC ≥ 0.5 and adjusted *q-value (FDR) < 0.05*) in response to hESC-RPE-sEV treatment, with 289 genes upregulated and 158 genes downregulated in treated RCS rats compared to untreated RCS rats. Additionally, we compared these findings with LE rat retinas unaffected by retinal degeneration. Of 447 DEGs, 394 (88%) between treatment and disease reversed expression significantly toward the healthy state after hESC-RPE-sEV treatment (**Supplement Figure A**). This highlights the therapeutic impact of hESC-RPE-sEVs on gene expression broadly associated with retinal degeneration in RCS rats. The Ingenuity Pathway Analysis (IPA) of the identified DEGs indicated that hESC-RPE-sEV treatment targeted genes associated with 145 Ingenuity Canonical Pathways (specific data not shown). There was a special focus related to “retina.”

The heatmap, emphasizing the top 65 DEGs associated with “signalizing pathways relevant to the retina” reveals that all but five genes displayed a reversal of expression towards healthy state levels following treatment across the three groups (**Figure 4-A**). Notably, several signaling pathways were significantly regulated, including those related to phototransduction and retinol metabolism. Among the DEGs related to phototransduction, *Cnga1*, *Slc24a1*, *Guca1n*, *Rho* and *Rcvm* were identified as the top 5 significantly upregulated genes after sEV treatment (**Figure 4-B and Supplement Figure B**). In the category of retinol metabolism-related genes, *Rdh8*, *Rdh11* and *Rdh12* were significantly upregulated after sEV treatment (**Figure 4-C and Supplement Figure C)**. Additionally, we observed significant upregulation of genes related to retinal lipid metabolism, involved in unsaturated fatty acids biosynthesis (*Elovl2* and *Elovl4*), sphingolipid metabolism (*Plpp2* and *Cerk*), and glycerophospholipid metabolism (*Dgke* and *Agpat5*) (**Figure 4-D**, **4-E**, **4-F and 4-G)**.

**Figure 4.**
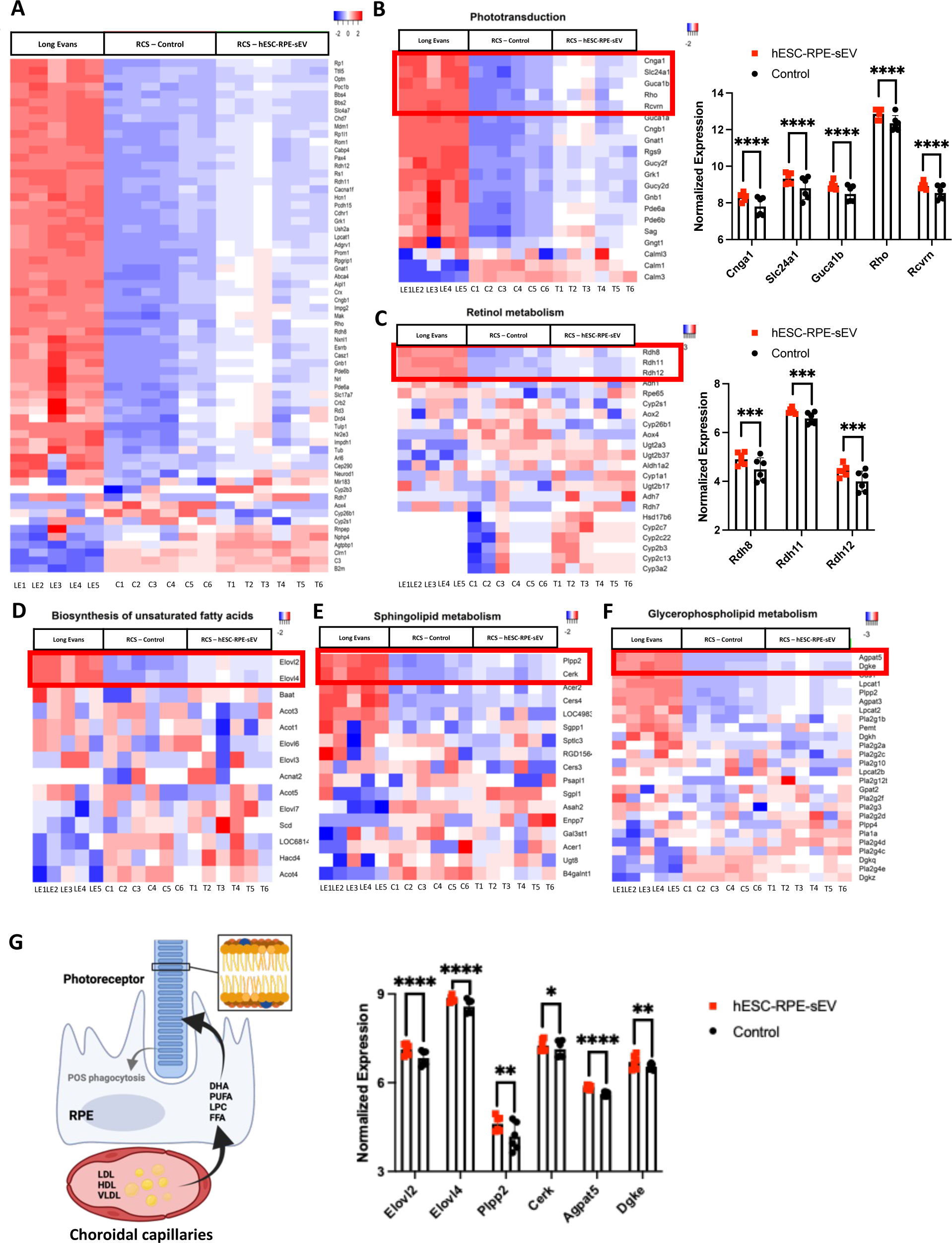
Transcriptomic analysis. Two consecutive intravitreal injection of hESC-RPE-sEV (∼ 5 x 10^8^/10 μl) (*n*=6) were administered to one eye of RCS Rats at P21 and P35 in comparison of untreated disease controls (*n*=6) and healthy retina controls in Long-Evans rat (*n*=5). **A**. The heatmap, emphasizing the top 65 DEGs associated with “signalizing pathways relevant to the retina,” reveals that all but five genes displayed a reversal of expression towards healthy state levels following treatment across the three groups. **B**. The heatmap illustrates 20 DEGs related to phototransduction, with *Cnga1*, *Slc24a1*, *Guca1n*, *Rho* and *Rcvm* as the top 5 significantly upregulated genes after sEV treatment. **C**. The heatmap illustrates 22 DEGs related to retinol metabolism, with *Rdh8*, *Rdh11* and *Rdh12* as the top 3 significantly upregulated genes after sEV treatment. **D-G** The heatmap reveals 14, 17, and 26 DEGs associated with unsaturated fatty acid biosynthesis (**D**), sphingolipid metabolism (**E**), and glycerophospholipid metabolism (**F**), respectively. *Elovl2* and *Elovl4*, *Plpp2* and *Cerk*, and *Dgke* and *Agpat5* emerge as the top upregulated genes following sEV treatment (**G**).

### hES-RPE-sEV treatment increase photoreceptor outer segment engulfment in RCS rat

Despite intrinsic *MerTK* mutation in RCS rats, POS engulfment significantly increased by 92.77% (*n*=3, *p-value 0.02*) after hESC-RPE-sEV treatments (**Figure 5-A**). Key genes involved in POS engulfment were reanalyzed from matched RCS retina’s transcriptomic analysis. While mutated *MerTK* expression remained unchanged, expressions of *Gas6*, *Adam9*, and *Pros8* were significantly upregulated after hESC-RPE-sEV treatments (**Figure 5-B**).

**Figure 5.**
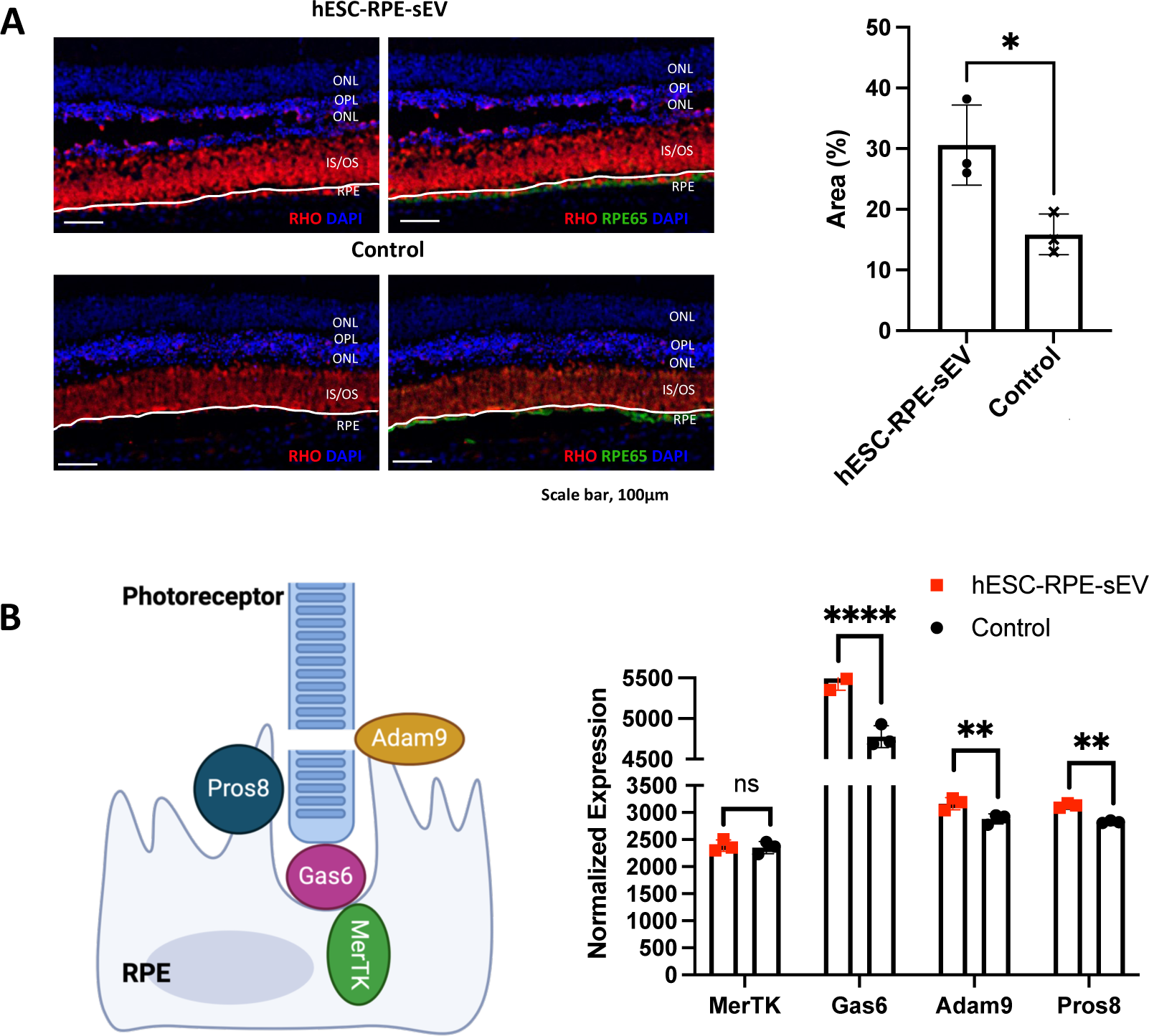
Photoreceptor outer segment phagocytosis assay. **A.** Representative immunofluorescence images of RHO (*red*) and RPE-65 (*green*) staining in RCS rat retinas at P49 with or without hESC-RPE-sEV treatments at P21 and P35 to assess prorector outer segment (POS) engulfment (*n*=3, *p-value 0.02*). **B**. Key genes involved in POS engulfment were reanalyzed from matched transcriptomic analysis. Scale bar, 100 μm. * *p*-value < 0.05, ** *p*-value < 0.01, **** *p*-value <0.0001.

## DISCUSSION

In our current study, we demonstrated that sEV secreted from fully differentiated RPE cells derived from hESC-RPE can effectively penetrate the full thickness of the retina local delivery via intravitreal administration. We first validated that the sEV fraction, rather than the sEV-depleted fraction of CCM from hESC-RPE (CCM minus sEV fraction), possesses the bioactive properties responsible for rescuing photoreceptors in immunocompromised RCS rats. This supported our hypothesis that sEV fraction is a major therapeutic component within RPE secretome. Additionally, our study results revealed that the xenogeneic intraocular administration of human cell-derived sEV in rodents did not induce immediate reactive retinal gliosis or acute immune responses. These findings collectively advocated for exploring the therapeutic potential of RPE-secreted sEV in models of retinal degeneration.

Testing in immunocompetent RCS rats, we observed that intravitreally administered hESC-RPE-sEV effectively rescued photoreceptors and restored their functionality. Furthermore, we observed a reduction in naturally elevated reactive retinal gliosis in RCS rats following treatment with hESC-RPE-sEV. RCS rats, characterized by photoreceptor cell death and retinal degeneration resulting from a mutation in the *MerTK* gene in RPE, have been well established as a valuable animal model for retinal degeneration.**19, 20** This mutation impairs the essential POS by RPE cells due to defective engulfment of POS to RPE, resulting in diminished POS clearance. This deficiency causes rapid photoreceptor cell death, thereby recapitulating a retinal disease condition where photoreceptor cell death is driven by dysfunctional RPE. Defective RPE phagocytosis, as seen in conditions like *MerTK* mutation, leads to retinal degeneration characterized by PR death and blindness in both human and animal models.**19, 21-23**

Additionally, our bulk RNA sequencing analysis, aimed at elucidating further the underlying mechanisms by which sEV released from healthy RPE cells rescue photoreceptor cells and restore their function in RCS rats, identified a total of 447 DEGs in response to hESC-RPE-sEV treatment, with 289 genes upregulated and 158 genes downregulated in treated RCS rats compared to untreated RCS rats. More importantly, 394 of the 427 DEGs (88%) between treatment and disease reversed expression significantly toward healthy state after hESC-RPE-sEV treatment. This supports that hESC-RPE-sEV treatment effectively reversed (“realigned”) the overall retinal transcriptomic changes observed in diseased retina of RCS rats when compared to healthy retina of Long-Evans rats. Based on the subgroup gene analysis focusing on relevant “retina pathways,” we identified the upregulation of multiple genes, including *Guca1n*, *Rho*, and *Rcvm*, essential for phototransduction, as well as *Rdh8*, *Rdh11*, and *Rdh12* essential for retinol metabolism crucial for the visual cycle following hESC-RPE-sEV treatment in RCS rats.

To our surprise, we observed upregulation of genes involved in essential retinal lipid metabolism, including unsaturated fatty acids biosynthesis (*Elovl2* and *Elovl4*), sphingolipid metabolism (*Plpp2* and *Cerk*), and glycerophospholipid metabolism (*Dgke* and *Agpat5*) after hESC-RPE-sEV treatment. Recent study by Mead et al. highlighted severe alterations in lipid metabolism in RCS rats, indicating that lipid metabolism is a potential therapeutic target for retinal degeneration.**23** The upregulation of genes associated with retinal lipid metabolism involving fatty acid synthesis, sphingolipid metabolism, and glycerophospholipid metabolism after hESC-RPE-sEV treatment is intriguing. The retina has substantial lipid demands like the central nervous system (CNS), necessitating either de novo synthesis of cholesterol or its acquisition from external sources like lipoproteins in the bloodstream and nutrition.**24-25** Both photoreceptor and RPE cells rely on tightly regulated lipid homeostasis. Their symbiotic relationship relies on the continuous incorporation and processing of each other’s membranes. In turn, PR cells regenerate stacked disks, piling lipid-bilayered structures on top of each other within the photoreceptor plasma membrane. This complex process necessitates the maintenance of membrane composition, including major lipid classes such as phospholipids, sterols, and free fatty acids (FFAs), at physiologically optimal levels for efficient phototransduction. Lipids also serve as an energy source and secondary signaling molecules. **25** Additionally, recent research has shown a downregulation of *Elovl2*, an enzyme involved in the elongation of long-chain polyunsaturated fatty acids, in the aged retina.**26** Therefore, it is particularly intriguing that *Elovl2* expression has been significantly upregulated following hESC-RPE-sEV treatment in RCS rat retinas. Furthermore, recent studies have revealed that the RPE secretes distinct EV from its apical side towards the photoreceptors and from its basal side towards the choroid.**27-29** Therefore, our study findings, alongside the enriched lipid-enriched composition of EVs and the emerging understanding of the role of RPE-secreted EVs, suggest a potentially substantial contribution to lipid homeostasis between the RPE and photoreceptor cells, thereby influencing overall retinal health. Further investigation is essential to fully comprehend the potential therapeutic implications of RPE-secreted EVs, especially in regulating lipid metabolism amidst the progression of retinal degenerative conditions.

We also observed that despite intrinsic *MerTK* mutations, hESC-RPE-sEV treatment increased POS engulfment. Additionally, we confirmed an upregulation of *Gas6*, *Adam9*, and *Pros8* genes, which encode key proteins involved in the process of POS engulfment, following sEV treatment. However, from the current dataset, it remains unclear whether hESC-RPE-sEV treatment compensated for the POS engulfment process or potentially activated alternative pathways for POS engulfment. While further investigation into the exact mechanisms involved in the restoration of RPE phagocytosis is warranted, our findings suggest an additional potential therapeutic benefit of hESC-RPE-sEV in restoring impaired RPE phagocytosis by supplying necessary bioactive molecules required for POS phagocytosis. Phagocytosis of shed POS by RPE is critical for maintaining retinal health and function. Each RPE cell, in contact with ∼30 photoreceptor cells is responsible for the phagocytosis and removal of the entire POS every 10-14 days.**30-33** In return, PR regenerate the POS in every 10-14 days, playing an essential role in phototransduction. Previous studies indicate that exposure to environmental factors such as excess iron (Fe) and cigarette smoke, as well as the natural aging process associated with age-related macular degeneration (AMD), can diminish RPE phagocytosis.**34-35** Whether caused by genetic mutations or age-related damage, impaired or reduced RPE phagocytosis leads to the accumulation of unphagocytosed debris, contributing to chronic inflammation. This disruption in the innate immune system balance exacerbates retinal damage.**36** This approach may serve as a protective biological pathway for photoreceptor cells, laying the groundwork for RPE-sEV therapy.

The findings from transcriptomic analyses collectively provide valuable insights into the therapeutic promise of RPE-secreted EVs in tackling a wider spectrum of retinal degenerative conditions. Earlier studies, reporting the therapeutic effect of intraocular EVs derived from various cell types, particularly stem cells, suggested that the primary mechanisms involved are immunomodulation and neuroprotection.**37-39** However, our results suggests that functional RPE cell recreated sEV may restore their functions to certain extent. This aligns with the current understanding that EVs participate in cell-to-cell communication by delivering a wide array of bioactive molecules, including proteins, lipids, and nucleic acids, thereby influencing gene expression in target cells.**1-4** Lessons from stem cell derived RPE cell transplantation studies also support that the paracrine actions of transplanted RPE cells are important therapeutic mechanism.**8-11**

In our current study, we observed that sEV-depleted CCM (CCM minus sEV fraction) failed to rescue photoreceptor cells, whereas sEV fraction effectively rescued photoreceptor cells and restored their function. Meanwhile, in our previous investigation, we demonstrated that the concentrated CCM of hESC-RPE, presumed to contain a moderately EV-enriched secretome fraction along with non-EV bound identified growth factors, comprises various neurotrophic growth factors secreted from RPE cells.**13** It is important to distinguish between near pure sEV fraction and a moderately EV-enriched fraction, which may contain other molecules such as growth factors, freely bound in CCM. Such a distinction is critical for identifying predominant bioactive molecules, responsible for observed effects and optimizing the formulation for improved efficacy and safety in clinical applications. Therefore, to advance intraocular EV therapeutics to the next stage, further studies must prioritize the identification of the active biological components within the RPE secretome, correlating them with the underlying mechanism of action. It is important to note that the various concentration of each fraction obtained during their preparation may affect the functional outcome in in vivo studies; therefore, studies with different doses may be needed to achieve precision.

Whether it involves the use of pure EVs or an EV-enriched fraction of the cell secretome, allogeneic treatment is likely in the context of human application. Immune reactions are relatively well-studied in allogeneic cell therapy, typically requiring immunosuppression. While there have been reports of no serious immune reactions observed in xenogeneic EV applications using animal models treated with human cell-derived EVs including our results, comprehensive studies on immune reactions after long-term allogeneic EV therapy are still lacking.**40-41** Therefore, additional long-term studies on immune reactions and toxicity are necessary for the translation of intraocular EV therapeutics.

It is noteworthy that intraocular EV therapy offers several additional potential advantages compared to direct stem cell implantation: (*i*) it can be administered locally via intravitreal injection, avoiding the invasive surgery, making it suitable for the early stages of retinal degeneration, (*ii*) it enables long-term therapy by allowing for multiple administrations, and (*iii*) a consistent potency threshold can be upheld from a replicating cell farm, whereas implanted stem cells may undergo aging over time.

Our study has limitations as our hypothesis was exclusively tested in RCS rats using a single dose of sEV solely secreted from hESC-derived RPE. Bulk RNA sequencing is limited to dissect transcriptomic changes in an individual cellular level given that hESC-RPE-sEV treatment broadly reversed transcriptomic changes within the retina. It is possible that other retina cell types such as microglia or macrophage could have been affected by the sEV treatment.

Further investigations employing various retinal degeneration models, as well as exploring different pluripotent stem cell-derived RPE such as hiPSC and implementing multiple dosing regimens will comprehensively assess the full potential of intraocular RPE -secreted EV therapeutics. Additionally, genomic analysis at the single-cell level will enhance the current understanding of the physiological role of sEV within retinal tissues.

In summary, our study represents the first, to the best of our knowledge, to demonstrate the therapeutic potential of RPE-secreted sEV, showing therapeutic efficacy in rescuing photoreceptors and their function by their ability to potentially restore RPE function and comprehensively align microenvironments within the retina. RPE-secreted sEV hold significant promise in supplementing RPE function by delivering a diverse array of molecules to address RPE dysfunction in various retinal diseases. To translate intraocular EV therapeutics into human applications, future studies should prioritize three key areas. Firstly, identifying the active biological components, which are likely to be multiple, aligning with various cells and pathways involved. This is crucial for establishing potency assays and scaling up the pipeline while maintaining consistency. Secondly, conducting long-term studies on immunologic and toxicity aspects following allogeneic or xenogeneic EV treatment is essential for understanding the safety and efficacy over extended durations. Lastly, the development of a sustained delivery system for intraocular EV application is imperative to ensure prolonged effectiveness.

## METHODS

### hESC-RPE Cell Differentiation, and Culture

The differentiation process of human embryonic stem cells (WA09, WiCell Research Institute) into RPE cells followed a previously outlined method.**42** In summary, hESC colonies underwent manual passage, were seeded onto Matrigel hESC-Qualified Matrix-coated tissue culture plates and underwent biweekly medium exchanges with serum free XVIVO-10 medium (Lonza #(BE) BP04-743Q) for around three months. Pigmented regions were manually isolated, expanded for two passages, and cryopreserved at 2-5 days post-seed as a cellular suspension using CryoStor10 cryopreservation medium (BioLife Solutions #210102). Thawed cells were expanded with 10uM Y-27632 (Tocris #1254) until the eighth passage, following a previously described protocol.**43-44** Cultures underwent testing every other month using the MycoAlert Mycoplasma Detection Kit (Lonza LT07-318) and consistently tested negative for mycoplasma throughout the study.

### hESC-RPE Pigmentation Assessment

When reaching confluence, representative fields of view were captured using bright field and phase contrast microscopy. Infinity Capture software was employed for the imaging process. The quantification of pigmented areas followed a previously outlined grading procedure.**13, 18**

### Immunocytochemistry and Confocal Fluorescence Microscopy of hESC-RPE Cells

hESC-RPE cells were cultured on sterile, matrigel-coated #1.5 coverslips (GG121.5PRE, Neuvitro) before fixation with 4% methanol-free formaldehyde in PBS for 20 minutes. Subsequently, cells were permeabilized with 0.1% Triton X-100 for 10 minutes, blocked with a mixture of 5% goat serum (Jackson ImmunoResearch) and 1% BSA (Thermo Fisher Scientific) in PBS for 30 minutes, and then subjected to overnight incubation at 4°C with the following primary antibodies: rabbit-anti-ZO1 (40-2200, Thermo Fisher Scientific) and mouse-anti-RPE65 (MAB5428, Millipore-Sigma). Coverslips underwent three PBS washes and were subsequently incubated with secondary antibodies in a blocking buffer for 1 hour at room temperature. The secondary antibodies used were AlexaFluor 594 AffiniPure goat anti-rabbit IgG (111585144, Jackson ImmunoResearch) and AlexaFluor 488 AffiniPure goat anti-mouse IgG (115545062, Jackson ImmunoResearch). Nuclei were stained for 10 minutes with Hoechst 33342 in PBS (2 μg/mL), followed by three PBS washes, mounting in ProLong Gold antifade, and imaging on an Olympus FV1000 Spectral Confocal with a PLAPON-SC 60X oil objective (NA: 1.40) and excitation laser lines at 405, 488, 559, and 635 nm. Single confocal planes extracted from z-stacks were generated using ImageJ-FIJI (NIH, USA).

### Preparation of Conditioned Cell Culture Medium

To generate conditioned cell culture medium (CCM) from hESC-RPE evaluation, RPE cells were enzymatically passaged using TrypLE Select (Gibco #12563011) per manufacturer’s instructions and seeded onto tissue culture plates coated with Matrigel hESC-Qualified Matrix (Corning #354277) at a density of 70,000 cells/cm2. The viability of the cells at the time of plating was determined by trypan blue exclusion, and cultures exceeding 95% viability were used for the study. At one day post-seed, cells were rinsed with DPBS (Gibco #14040141) and fed with 4mL XVIVO-10 medium (Lonza #(BE)BP04-743Q) per well within a 6-well tissue culture plate. The medium was exchanged twice per week and was supplemented with 10mM Y-27632 (Tocris #1254) for 10-15 days until the RPE cells attained cuboidal morphology. From day 28 to 39 post-seed, CCM was collected from four wells of a 6-well plate using a serological pipet, combined, and immediately frozen in a 50mL Falcon centrifuge tube at -80°C for future use. Four replicate plates were used at each collection time point. The incubation period during which cells conditioned the medium ranged from 3-4 days at 37°C, 5% CO2.

### Small Extracellular Vesicle Recovery

To recover sEV, ExoDisc® (LabSpinner, Ulsan, South Korea), a microfluidic tangential flow filtration method-based equipment, was utilized according to the manufacturer’s instructions.**44** In brief, 3 mL of hESC-RPE CCM was processed on an ExoDisc® using the bench-top operating machine (OPR-1000, LabSpinner™, South Korea). Purified sEV were obtained from the collection chamber using 100 μL of PBS and along with the sEV-depleted part of CCM promptly stored at −80 °C for future use within 2-4 weeks.

### Biophysical Particle Analysis

To determine the concentration and size distribution of the recovered sEV, NanoSight (Malvern Instruments Ltd., UK) was employed following the manufacturer’s protocol. In a nutshell, for NanoSight (NS) analysis, the samples were appropriately diluted to achieve an optimal detection concentration of 108 particles/mL. A constant flow speed of 25 was maintained using an automated syringe pump, and five videos were recorded with a camera level set to 14. The collected data were analyzed using NTA software 4.3 with a detection threshold of 7, and adjustments were made based on the dilution factor.

### Transmission Electron Microscopy

To visualize sEV, negative-stained transmission electron microscopy (TEM) was conducted using a JEOL JEM-2100 microscope equipped with a Gatan OneView IS camera as previously described.**44**

### Single-Particle Interferometric Reflectance Imaging Sensing: ExoView Analysis

The ExoView R100 system along with the ExoView Human Tetraspanin Kit from NanoView Biosciences, USA, was employed for Single Particle Interferometric Reflectance Imaging Sensing (SP-IRIS) as previously described.**44 A** from the ExoView Human Tetraspanin Kit (NanoView Biosciences, USA). Immunocapture antibodies (anti-CD9 CF488, anti-CD81 CF555, and anti-CD63 CF647), pre-diluted to a 1:500 ratio in solution A, were utilized.

### Bead-Based Multiplex Flow Cytometry Assay (MACSPlex) Analysis

The assessment of expanded surface protein markers on sEV was carried out using a MACSPlex human Exosome kit (Miltenyi Biotec, Bergisch-Gladbach, Germany) following the manufacturer’s protocol.**44** Thirty seven distinct surface marker antibodies (CD1c, CD2, CD3, CD4, CD8, CD9, CD11c, CD14, CD19, CD20, CD24, CD25, CD29, CD31, CD40, CD41b, CD42a, CD44, CD45, CD49e, CD56, CD62p, CD63, CD69, CD81, CD86, CD105, CD133.1, CD142, CD146, CD209, CD326, HLA-ABC, HLA-DR DP DQ, MCSP, ROR1, and SSEA-4) were utilized to capture sEV simultaneously.

### Protein concentration analysis and Western Blot

The isolated sEV underwent lysis by combining equal parts of the sample and 2x RIPA buffer containing protease inhibitors. The protein concentrations of the lysed sEV preparations were determined using the Pierce™ BCA Protein Assay Kit (Thermo Fisher Scientific, USA), according to the provided guidelines. For the Western blot analysis, the isolated protein was further concentrated using Microcon® Centrifugal Filter Devices 10KD (12 minutes at 4°C and 14,000g). A Bradford protein assay was performed again and for gel loading, 10 µg of protein was utilized, mixed either with non-reducing 4X SDS Lumina buffer or reducing 4X SDS Laemmli buffer supplemented with Beta-mercaptoethanol. As controls, 10 µg of protein lysate from HEK293 cells was loaded alongside the experimental samples. The primary antibodies used were: CD9 (HI9a, 1:1000), CD63 (BD Bioscience #556019, 1:1000), Calnexin (C5C9, 1:200), Fibronectin (EP5, 1:200), HSC70/HSP70 (W27, 1:200) and CD81 (D3N2D, 1:1000). The secondary antibodies used were: anti-mouse 800CW (1:10000), anti-mouse 680RD (1:10000), and anti-rabbit 800CW (1:10000).

### Animals

The treatment and handling of rats was adhered to the regulations and guidelines set forth by the Institutional Animal Care and Use Committee (IACUC) of the University of Southern California (USC), the National Institutes of Health (NIH) Guidelines for the Care and Use of Laboratory Animals and the Association for Research in Vision and Ophthalmology (ARVO) Statement for the Use of Animals in Ophthalmic and Vision Research. Immunocompetent or immunocompromised Royal College of Surgeons (RCS) rats, characterized by retinal degeneration due to the *MerTK* mutation and Long-Evans rats, as no retinal degeneration control, were employed. **19** C57BL/6J mice were also utilized.

### Intravitreal Injection of hESC-RPE-sEV

RCS rats were subjected to intravitreal (ITV) injections of hESC-RPE-sEV (∼ 5 x 10^8^/10 μl) at the age of 21 days (P21) and 35 days (P35). The left eye (OS), conjunctival sac, and eyelids were cleaned with a 5% povidone-iodine solution, and topical anesthesia was administered. Utilizing fine forceps, both superior and inferior eyelids were retracted, and a temporal superior quadrant sclerotomy, along with a corneal paracentesis 180° apart, was performed using a 30G sterile needle. The IVT injection involved delivering 10 μl of the treatment under a surgical microscope, employing a 33G blunt needle attached to a 25 μL syringe (Hamilton Syringe, Hamilton Company, Switzerland). Wild type mice received single IVT injection of hESC-RPE-sEV (∼ 5 x 10^7^/1 μl) to assess any acute immune reaction. All the procedures were executed under sterile conditions.

### In vivo imaging

Optical coherence tomography (OCT) imaging was carried out utilizing a commercially available cSLO (SPECTRALIS HRA+OCT, Heidelberg, Germany) for all animals. To facilitate imaging, pupils were dilated with 2.5% phenylephrine hydrochloride and 1% tropicamide. The imaging sessions took place on P21 (baseline), P35, and P49. A high-resolution b-scan was captured from the temporal side of the optic nerve for both eyes and subsequently, the outer nuclear layer (ONL) thickness was measured using ImageJ.

### Electroretinogram

The assessment of ERG was conducted at P21 (baseline) and P49. Both scotopic testing involved flash stimuli with intensities ranging from 0.1-25 candela (cd) and photopic testing with with intensities from 0.01-25 cd using the HMsERG Rodent System from OcuScience, NV were conducted as previously described. **13** The raw data from ERG assessments were collected and analyzed.

### 2.15 Immunocytochemistry and Confocal Fluorescence Microscopy of Retina Tissue

The rats were euthanized at P49, and their eyes were harvested for either histological or transcriptomics analysis. For histological analysis, eyes were fixed in 4% PFA, embedded in an optimal cutting temperature medium (OCT; Sakura, Japan), frozen in liquid nitrogen, and stored at −80 °C for future use. A Microm HM 550 cryostat was used to obtain 8 μm thick frozen sections. The sections were permeabilized with 0.1% Triton X-100 and blocked with a mixture of 10% goat serum (Jackson ImmunoResearch) in PBS for 1 hour, and then subjected to overnight incubation at 4°C with the primary antibodies: rabbit anti-GFAP (Z0334, DAKO), mouse anti-Rhodopsin (AB5417, Abcam), and rabbit anti-RPE65 (AB231782, Abcam) antibodies. The secondary antibodies used were anti-mouse Alexa Fluor 594 (A11005, Invitrogen), anti-rabbit Alexa Fluor 488 (A11034, Invitrogen), and 4′6′-diamino-2-phenylindole (DAPI, H-1500, Vector Laboratories) for nuclei staining. Stained frozen sections were examined using a ZEISS Observer Z1 confocal microscope.

### Intraocular distribution of hESC-RPE-sEV

To fluorescently label the sEV, MemGLowTM 640 (# MG04-02, Cytoskeleton, Inc, USA) was utilized following the manufacturer’s instructions. Briefly, 0.5 μL of 20 μM MemGLowTM stock was diluted in 100 μL of recovered sEV-hESC-RPE in PBS. The labeled vesicles were then intravitreally injected into Long-Evans rat eyes using the previously described method. Animals were euthanized either after 1 or 7 days post-injection, and the harvested eyes were sectioned and stained with DAPI as previously outlined. The frozen sections were also stained with anti-GFAP (Z0334, DAKO) and anti-rabbit Alexa Fluor 488 (A11034, Invitrogen) to evaluate the presence of reactive gliosis after the IVT injection of the sEV-hESC-RPE. The stained sections were examined using a ZEISS Observer Z1 confocal microscope.

### Assessing retinal acute immune reaction from hESC-RPE-sEV

To further assess the any retinal acute immune reaction from ed hESC-RPE-sEV treatment, C57BL/6J mice received IVT injection of either hESC-RPE-sEV (*n*=6), Lipopolysaccharides (LPS) (*n*=3) or PBS (*n*=3). Retinas (3/ group) were harvested 1 day after the hESC-RPE-sEV (∼ 5 x 10^7^/μl), LPS (1 ng/μL), and PBS injection and 7 days after the hESC-RPE-sEV injection. Total RNA was purified using RNeasy columns (Qiagen) and included an on-column genomic DNA digestion step using RNase-free DNase (Qiagen). Reverse transcription was performed on 1 ng of total RNA using a commercial kit (High Capacity RNA-to-cDNA Kit; Applied Biosystems) and Real-Time PCR amplification (CFX Opus 96 Real-Time PCR Instrument, Biorad) was performed using TaqMan Universal PCR Master Mix (Applied Biosystems) and the following TaqMan probe/primers (Applied Biosystems): TNF-α (Mm00443258_m1), IL-4 (Mm00445259_m1), IL-6 (Mm00446190_m1), IL-13 (Mm00434204_m1), IFN-γ (Mm01168134_m1) and GAPDH (Mm99999915_g1). All Ct values for TNF-α, lL-4, IL-6, IL-13, and IFN-γ were normalized to the reference gene GAPDH.

### Image analysis

Image analysis was conducted utilizing ImageJ software. To quantify GFAP expression, the confocal-acquired images were converted into 8-bit images, followed by a histogram analysis, normalized by DAPI intensity. For the assessment of photoreceptor engulfing/phagocytosis, the confocal-acquired images were subjected to color-thresholding to identify yellow-hued (green-red overlapping) areas, facilitating the measurement of Rhodopsin and RPE65 colocalization areas.**45**

### Transcriptomic analysis

The harvested retinal tissue (hESC-RPE-sEV treated RCS retinas, *n*=6; untreated RCS retinas, *n*=6; Long Evans retinas, *n*=5) was sent to Azenta Life Sciences (South Plainfield, NJ) for bulk RNA sequencing analysis. Sequencing was performed on an Illumina NovaSeq 6000 instrument with paired-end 150bp reads. Sequence reads were trimmed to remove possible adapter sequences and nucleotides with poor quality using Trimmomatic v.0.36. The trimmed reads were mapped to the Rattus norvegicus Rnor6.0 reference genome available on ENSEMBL using the STAR aligner v.2.5.2b. Unique gene hit counts were calculated by using featureCounts from the Subread package v.1.5.2. Read-count normalization and differentially expressed (DE) analyses were performed using the edgeR package from Bioconductor. Expression values quantile normalized with the voom function were analyzed for differential expression using the standard functions of the limma package. Moderate t-test p-values were adjusted for multiple testing using the false discovery rate (FDR) method and FDR (q-value)<0.05 was used to filter significant differences. Genes differentially expressed between treatment and disease samples, reversing to healthy expression levels after treatment (DisRev), were identified by collapsing healthy and treatment groups into a single group, and contrasting it to disease samples. Since the *t*-statistic considers the difference between group averages relative to within-group variation, a strong t-statistic would indicate both strong difference between disease samples versus the other groups, and similar expression levels in healthy and treatment groups.

### Pathway analysis

Gene Set Enrichment Analysis (GSEA), identifying sets of genes sharing the same functionality (GO, KEGG, Reactome pathways), overrepresented among the differentially expressed genes, was performed with specialized packages from Bioconductor such as fgsea, ReactomePA and viewPathway. Each pathway GSEA edge set was used as the basis of selection for the genes included in the heatmaps showing the pathway activity variation among the three types of samples (Healthy, Disease, Treatment) Ingenuity Pathway Analysis (IPA, QIAGEN, Redwood City CA, https://www.qiagenbioinformatics.com/products/ingenuitypathway-analysis) was used for further discovery and interactive exploration of significantly impacted static and causal gene networks, pathways, disease, upstream regulators, and regulatory effects.

### Statistical Analysis

Unless otherwise described, all values were reported as mean ± standard error. GraphPad Prism was used for the statistical analysis and graph plotting. Statistical differences were measured with Student’s t-test or Analysis of Variance (ANOVA) that was appropriate for each analysis. For small sample size and skewed data values robust testing was performed using Mann-Whitney and Kruskal-Wallis nonparametric test alternatives. The significance level was a priori predetermined for all comparisons with a threshold set at *p* <0.05.

## Acknowledgments

The authors thank Extracellular Vesicle Core at Children’s Hospital Los Angeles and Paolo Neviani, PhD, for their technical assistance with NanoSight and ExoView. Dr. Allen and Charlotte Ginsburg Institute for Biomedical Therapeutics (GF1000134); Dennis and Michele Slivinski (GF1000135); and P30EY029220

